# Quantitative Single-Molecule Imaging with Statistical Machine Learning

**DOI:** 10.1101/2021.07.30.454455

**Authors:** Artittaya Boonkird, Daniel F. Nino, Joshua N. Milstein

**Affiliations:** Department of Chemical and Physical Sciences, University of Toronto Mississauga, Mississauga, Ontario L5L 1C6, Canada; Department of Physics, University of Toronto, Toronto, Ontario M5S 1A7, Canada

## Abstract

Single-molecule localization microscopy (SMLM) is a super-resolution technique capable of rendering nanometer scale images of cellular structures. Recently, much effort has gone into developing SMLM into a quantitative method capable of determining the abundance and stoichiometry of macromolecular complexes. These methods often require knowledge of the complex photophysical properties of photoswitchable flourophores. We previously developed a simpler method built upon the observation that most photswitchable fluorophores emit an exponentially distributed number of blinks before photobleaching, but its utility was limited by the need to calibrate for the blinking distribution. Here we extend this method by incorporating a machine learning technique known as Expectation-Maximization (EM) and apply it to a statistical mixture model of monomers, dimers and trimers. We show that the protomer fractions and the underlying single-fluorophore blinking distributions can be inferred, simultaneously, from SMLM datasets, obviating the need for an additional calibration and greatly expanding the applicability of this technique. To illustrate the utility of our approach, we benchmark the method on both simulated datasets and experimental datasets assembled from dSTORM images of Alexa-647 labeled DNA nanostructures.

---

In recent years, a variety of single-molecule imaging methods have been developed that are capable of resolving sub-diffraction limited, nanometer scale cellular features. These techniques include (d)STORM [1–3], (f)PALM [4, 5], and DNA PAINT [6], among others, and are collectively referred to as single-molecule localization microscopy (SMLM). SMLM relies upon harnessing the stochastic blinking of fluorescent labels to sparsify a fluorescent signal. The coordinates of each labeled molecule can then be precisely determined and, from these localizations, a detailed image rendering can be constructed.

In addition to generating super-resolved images, the localization datasets acquired by SMLM contain the necessary information for counting single-molecules. If interpreted correctly, these datasets would reveal quantitative properties of macromolecular complexes such as their abundance, stoichiometry or oligomerization state. An active effort to advance quantitative SMLM (qSMLM) by developing methods to accurately extract this information is still in its infancy [7–20]. These methods attempt to correct for counting artifacts introduced by the stochastic blinking required for super-resolved imaging. They primarily rely upon translating either the number of localization events [9, 11, 15, 21] or the number of blinks [8, 10, 13, 17] into an accurate molecular count by incorporating additional information on the emission properties of the fluorophores, such as the ON and OFF times or rates [14, 19].

Utilizing the empirical fact that many photswitchable fluorophores emit an exponentially distributed number of blinks before photobleaching, we previously developed a statistical model for molecular counting with SMLM [17, 18]. The advantage of this approach is that one only needs to calibrate for the expected number of blinks λ defining the blink distribution for a particular fluorphore. Unfortunately, it is not always practical to perform this calibration, which limits the utility of the method. For example, we have shown that organic dyes can display an acute sensitivity to their environment, requiring repeated calibration from sample to sample within the cellular milieu under investigation [18].

In this manuscript, we show how the blinking distribution can be inferred directly from SMLM datasets by applying a machine learning algorithm known as Expectation-Maximization to a statistical mixture model of an ensemble of fluorescently labeled, macromolecular complexes. This advance obviates the need for a separate calibration of λ, so greatly expands the range of experimental conditions under which qSMLM based molecular counting can be performed. We first present the theory behind the learning algorithm, generalize it to account for labeling inefficiency, and then illustrate the accuracy of the method on both simulated datasets and experimental datasets assembled from dSTORM images of Alexa-647 labeled DNA nanostructures.

For *N* fluorophores that emit *B* blinks, the exponential, single fluorophore blink distribution generalizes to a negative-binomial distribution of the form

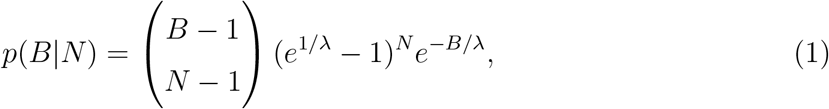

where we assume that each fluorophore detected emits at least one blink (*B* ≥ *N*) [17, 18]. Now consider a mixture of macromolecular complexes built from a maximum of *h* monomeric subunits, with each monomer labeled by a single, photoswitchable fluorphore. The probability of observing *B* blinks from this ensemble is given by the weighted sum of negative binomial distributions:

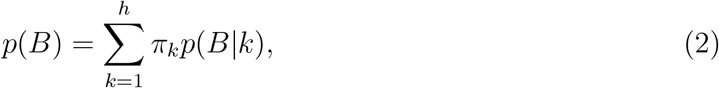

with weights *π_k_* = [*π*_1_, *π*_2_ … *π_h_*] defining the fraction of monomers, dimers, etc., within the ensemble. Since the number of blinks *B* is the experimental observable, Eq. 2 defines a mixture model for the probability *p*(*B*). It turns out that the unobserved, or latent, parameters λ and weights *π_k_* can all be estimated from the experimental data via Expectation-Maximization.

The Expectation-Maximization (EM) algorithm is a popular machine learning algorithm for determining latent variables within a statistical model [22]. The EM-algorithm iterates between two modes, until convergence, to perform a maximum likelihood estimation. For our purposes, these are composed of an E-step to estimate the values of the mixture weights *π_k_*, and an M-step that estimates λ by maximizing the likelihood of the model.

For each complex we image, up to *N_T_* in total, one might observe *B*^(1)^, *B*^(2)^…, *B*^(*N_T_*)^ blinks. For a given value of the latent variable λ, we define the posterior probabilities, or expectations, 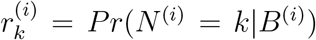 as the probability that complex *i* has *N*^(*i*)^ = *k* monomeric subunits given that it emitted *B*^(*i*)^ blinks. The expectations may be written as follows:

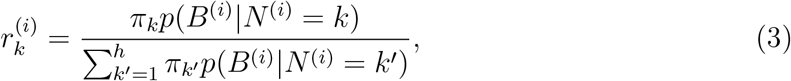

which follows from Bayes’ theorem. The weights *π_k_* are then simply estimated as 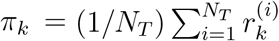, which defines the E-step of the algorithm.

Next, for the M-step of the algorithm, we need to compute the maximum likelihood estimate of λ. Toward this end, we define the log-likelihood function for our mixture model as

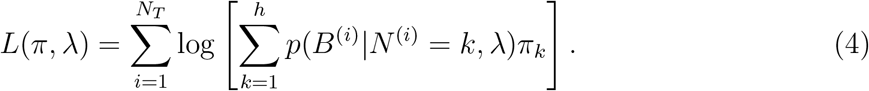

The maximum likelihood estimate for λ can be calculated by maximizing the log-likelihood (i.e., *dL*/*d*λ = 0). From Eq. 3, this may be written in terms of the expectations 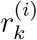 found in the previous E-step

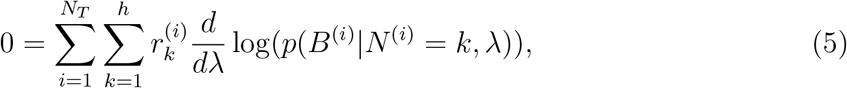

which may be solved for λ after substitution of Eq. 1:

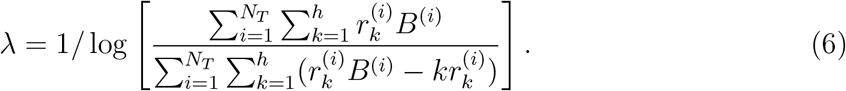

We have so far assumed a perfect one-to-one labeling of each monomer. To account for imperfect labeling, consider a protein complex with *h* subunits. If each subunit may accommodate at most 1 active fluorophore, the probability of having *N* active labels on the complex is given by a binomial distribution:

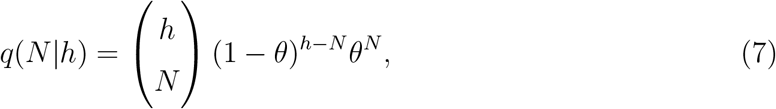

where *θ* is defined as the labeling efficiency. For covalent technologies that label proteins with organic dyes, such as SNAP-tags, *θ* can usually be obtained through appropriate control experiments, with values often exceeding ~ 0.9 [23]. Fluorescent proteins, on the other hand, may be directly co-expressed with a protein of interest, but *θ* must now account for incomplete maturation and the photoconversion efficiency. Fortunately, for many fluorescent proteins, *θ* may be measured a priori and appears to be a fairly robust property of the fluorophore [24–26].

Now let us incorporate the labeling efficiency into our mixture model of Eq. 2, which may be rewritten as

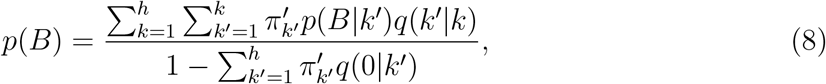

where the normalization term in the denominator arises because *p*(*B*|*N*) is defined for *N* ≥ 1, whereas *q*(*N|h*) spans *N* ≥ 0. Equation 8 is equivalent to our original mixture model, Eq. 2, with the weights of the latter scaled as follows:

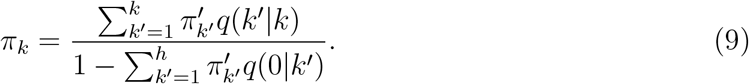

To account for fluorophore labeling inefficiencies, we simply need to perform an EM algorithm on the original mixture model to obtain the weights [*π*_1_, *π*_2_ … *π_h_*], then invert Eq. 9 to obtain the corrected fraction of monomers, dimers, etc. (i.e., 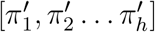). For *h* = 2, a mixture of monomers and dimers, the scaling is

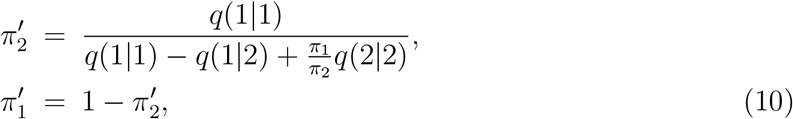

and for *h* = 3, a mixture of monomers, dimers and trimers, the scaling is

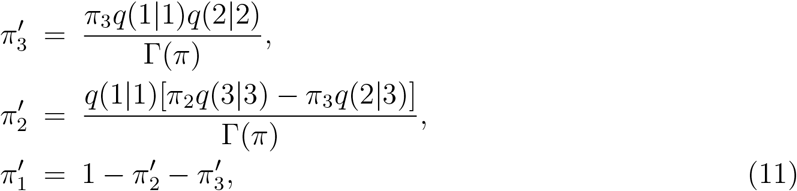

where,

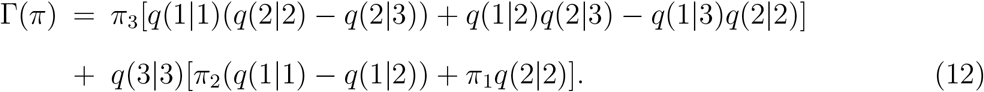

To assemble a control against which to benchmark the performance of this method, we acquired a series of dSTORM images of single Alexa-647 dyes centred within a DNA origami grid (to mitigate surface effects) and sparsely bound to a glass coverslip (see Supplemental Materials (SM) [27]). The number of emitted blinks from 1,095 individual dyes were first extracted from the image stacks. The blinking statistics were well described by an exponential and the characteristic number of blinks was found to be λ = 3.6 ± 0.2. Single-dye blinks were then assembled into subpopulations of monomers, dimers, and trimers, of relative target weights *π*_1_, *π*_2_, and *π*_3_, respectively. For each mixture, 1000 protomers were selected (each assembled from dyes chosen at random, but allowing for repetition) and the analysis was repeated 50 times to estimate both the mean and the uncertainty.

Figure 1 displays the results of this experimental benchmark. For the purely monomeric case (Fig. 1A (i)), although over-parameterized, both the monomer/dimer (M/D) and the monomer/dimer/trimer (M/D/T) model predict the majority of the population as monomers, while admitting a small fraction of higher order protomers. Note, a simple one-state monomer model is not shown since *π*_1_ = 1 by default. In Fig. 1A (ii), for a 0.8/0.2/0 mix (M/D/T), we find that both models accurately predict the monomeric and dimeric fractions. Once again, this is despite the over-parameterization of the M/D/T model, which tends to also predict a small trimeric fraction. In Figs. 1A (iii) and (iv), we challenged the algorithm to a target population at ratios 0.3/0.35/0.35 and 0.4/0.4/0.2, respectively. In both cases, the under-parameterized M/D model fails by predicting an abundance of dimers, while the M/D/T model accurately estimates each population.

**FIG. 1:**
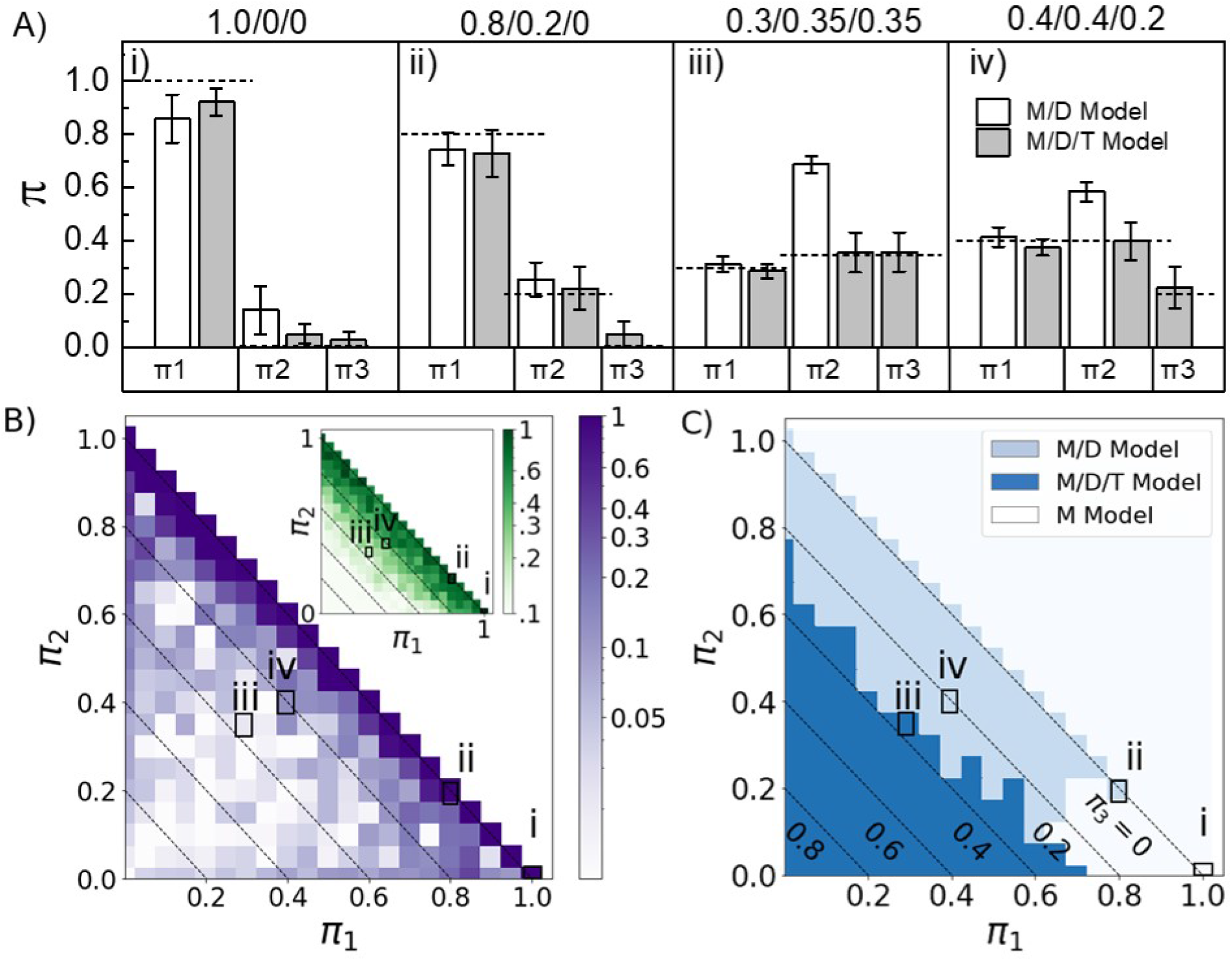
Performance of the algorithm on dSTORM Alexa-647 data (λ = 3.6±0.2, sample size=1,000 protomers, 50 replicates). A) Mean estimates and uncertainty of monomer (*π*_1_), dimer (*π*_2_) and trimer (*π*_3_) fractions for M/D/T populations: i) 1.0/0/0, ii) 0.2/0.8/0, iii) 0.4/0.4/0.2 and iv) 0.30/0.35/0.35. B) Mean relative error in *π*_3_ for the M/D/T model (insert shows the CV). Results for M/D/T models shown in Figs. A (i-iv) are indicated. C) Model selection based on AIC score (results for Figs. A (i-iv) are indicated).

In Fig. 1E, we show the mean relative error in predicting the dimeric fraction *π*_3_ by the M/D/T model over a range of protomer fractions. Because *π*_1_ + *π*_2_ + *π*_3_ = 1, we can specify all three weights in this two-dimensional heat-map, where the diagonal lines represent increments of the trimeric fraction *π*_3_. Indicated within the figure are the corresponding values for the M/D/T models shown in Figs. 1A (i-iv). We see that, on average, the algorithm is able to learn *π*_3_ to within roughly 10% of the true value across the majority of the parameter space. In the insert to Fig. 1B, we display the associated coefficient of variation (CV), which is the standard deviation normalized to the mean. The CV quantifies the relative uncertainty in the prediction of the trimeric fraction *π*_3_ for an individual measurement. As expected, the CV grows substantially as *π*_3_ → 0. For the two examples of a system containing a trimeric population shown here (iii and iv), 〈*π*_3_〉 = 0.36 and 0.23 with a CV of 0.21 and 0.35, respectively.

In many stoichiometric measurements, the actual degree of oligimerization is unknown. In these situations, the Akaike information criterion (AIC) may serve as a guide to selecting the appropriate probabilistic model for the data [28]. The AIC uses the maximum likelihood estimate of a model as a measure of fit, while penalizing the fit for each additional independent variable within the model. It is defined as follows:

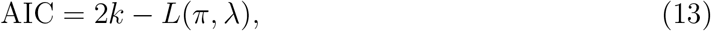

where *k* is the number of independent variables and *L*(*π*, λ) is the likelihood as given by Eq. 4. Note, while the variables here are the weights *π*_1_, *π*_2_… *π_h_* and λ, since the weights sum to 1, *k* = *h* and not *h* + 1.

The higher the AIC score, the better the model fits the available data. In Fig. 1C, we calculated the AIC scores across all values of *π*_1_, *π*_2_, and *π*_3_, then indicate the ranges in which the AIC score predicts either the M/D, M/D/T, or a purely monomeric model. The AIC prediction for the cases illustrated in Figs 1A (i-iv) are explicitly labeled within the figure. The AIC predicts the appropriate model for fractions (i-iii); however, for the mixture with the smaller trimeric fraction (iv), the AIC inaccurately selects for the M/D model. This is despite the fact that the M/D/T model accurately predicts the protomeric fractions. Like any predictive method, the AIC score is data limited, and it is accurate over a larger range of parameters with increasing sample size or decreasing λ (see SM [27]). So, in practice, while the AIC score may be used to support a particular model, it is not a definitive selection criterion.

We next consider the effects of imperfect labeling. Clusters are again assembled from the experimentally acquired dSTORM localization data of single Alexa-647 dyes, but this time we remove dyes at random dependent upon the labeling efficiency *θ*. The analysis is now repeated 200 times for each mixture to generate a convergent mean and uncertainty in the estimate. In Fig. 2, we consider the M/D/T population fractions 0.3/0.35/0.35 and 0.4/0.4/0.2. The algorithmic accuracy at predicting the mean of the fractions *π*_1_, *π*_2_ and *π*_3_ converges to within 20% of the true fraction down to a labeling efficiency of *θ* = 0.8, and is often well below that. Below *θ* = 0.8, the algorithm begins to struggle with the 0.3/0.35/0.35 population, although stays within 20% for the 0.4/0.4/0.2 population. The uncertainty of any single measurement is also strongly affected by *θ*. For the smallest trimeric fraction considered here, *π*_3_ = 0.2, the CV is already ~ 0.4 for perfect labeling increasing to ~ 0.7 at *θ* = 0.7 (Fig. 2C). However, the larger trimeric fraction, *π*_3_ = 0.35, displays a CV that sharply grows from ~ 0.3 when perfectly labeled to ~ 0.7 at *θ* = 0.7 (Fig. 2B). As an aside, the AIC consistently predicts the incorrect model at *θ* = 0. 9 or below.

**FIG. 2:**
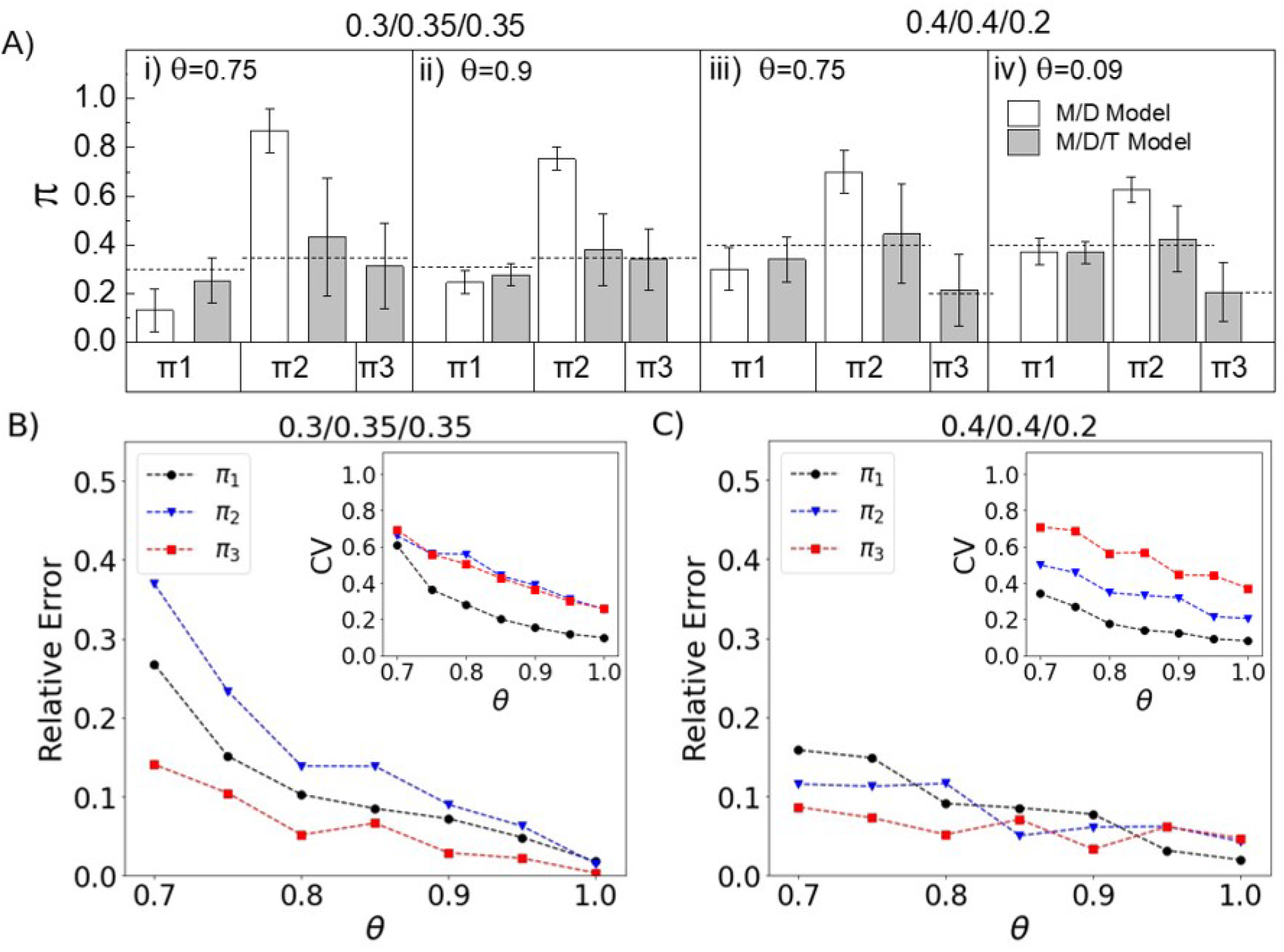
Effects of imperfect labeling on dSTORM Alexa-647 data (200 replicates, 1000 protomers each). A) Performance of the M/D and M/D/T models at predicting a population of 0.3/0.35/0.35 or 0.4/0.4/0.2 for labeling efficiencies of *θ* = 0.9 and 0.75. Relative error and CV (insert) as a function of the labeling efficiency for B) 0.3/0.35/0.35 and C) 0.4/0.4/0.2 populations.

Finally, we wish to understand how λ, which describes a chosen fluorophore’s propensity for blinking, affects the performance of this method. While the photoswitching of Alexa-647 can be tuned to a certain degree, we found it easier to increase λ than to decrease it much below ~ 4, so we instead resort to a series of numerical simulations (see SM [27]). In Fig. 3, we simulated the photoswitching of an M/D/T population at various fractions *π*_1_, *π*_2_ and *π*_3_ for λ = 2, 4 and 8. For each set of parameters, we performed 50 simulations of a population composed of 1000 protomers in total. The mean relative error in the estimates of *π*_3_ are shown in the figures, while the inserts display the relative uncertainty (CV) in these estimates. Note, for λ = 4 the results are similar to our experimental results (Fig. 1B) with the estimates falling within ~ 10% of the target values across most of the parameter space. This helps support any inferences obtained through these simple simulations. We find that decreasing (increasing) λ decreases (increases) the errors both in the estimates of *π*_3_ as well as the uncertainty in those estimates. While results for only the trimeric fraction *π*_3_ are shown, this trend holds for the monomeric and dimeric fractions *π*_1_ and *π*_2_.

**FIG. 3:**
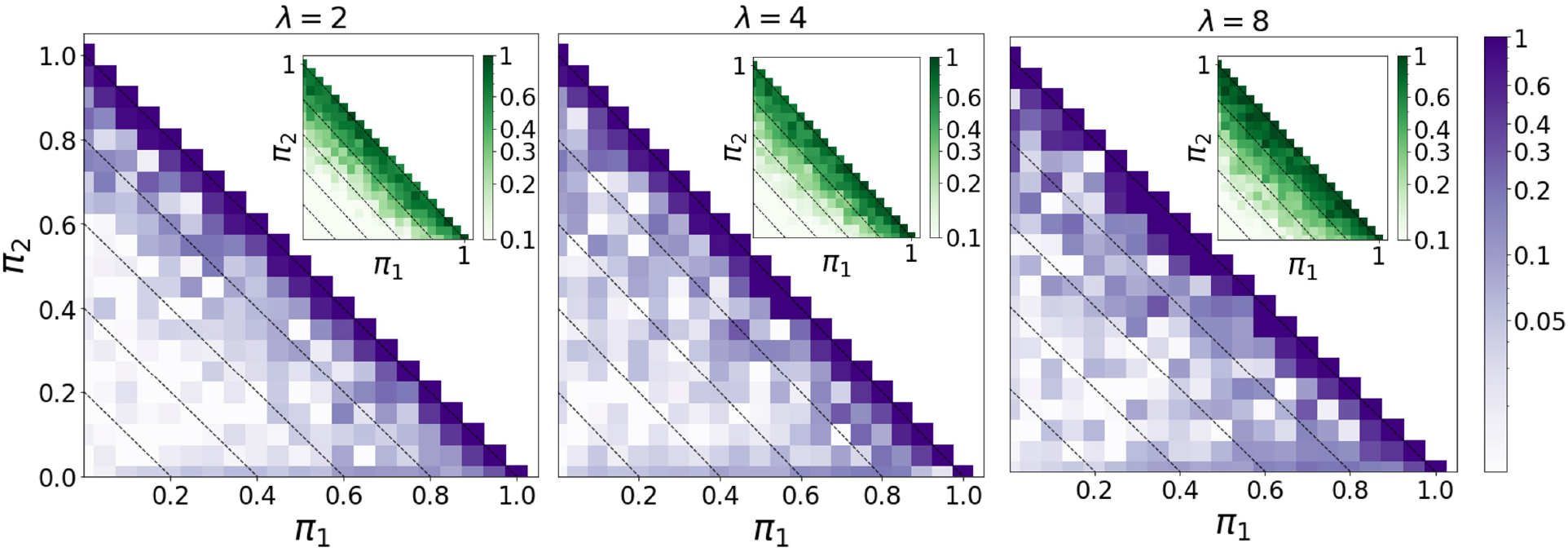
Effect of tuning λ on the algorithmic accuracy within the M/D/T model (simulations of 50 replicates, 1000 protomers each). From left to right, the figures display the mean relative error in predicting the dimeric fraction *π*_3_ for λ =2,4 and 8. Inserts show the corresponding uncertainty (CV).

In Fig. 4, we incorporate the added effects of an imperfect labeling efficiency. Here we only consider the M/D/T model for mixtures of 0.3/0.35/0.35 or 0.4/0.4/0.2, and contrast λ = 4 and λ = 2. For each set of parameters, we performed 200 simulations of a population composed of 1000 protomers in total to obtain converged values for the means and uncertainties of the estimates. Once again, a reduced labeling efficiency degrades the predictions of *π*_1_, *π*_2_ and *π*_3_, but has its most pronounced effects on the uncertainty in the measurement. We find that while a smaller λ may reduce the uncertainty in a well labeled system, these gains are modest and unable to compensate for the growing uncertainty that results with decreased labeling efficiency.

**FIG. 4:**
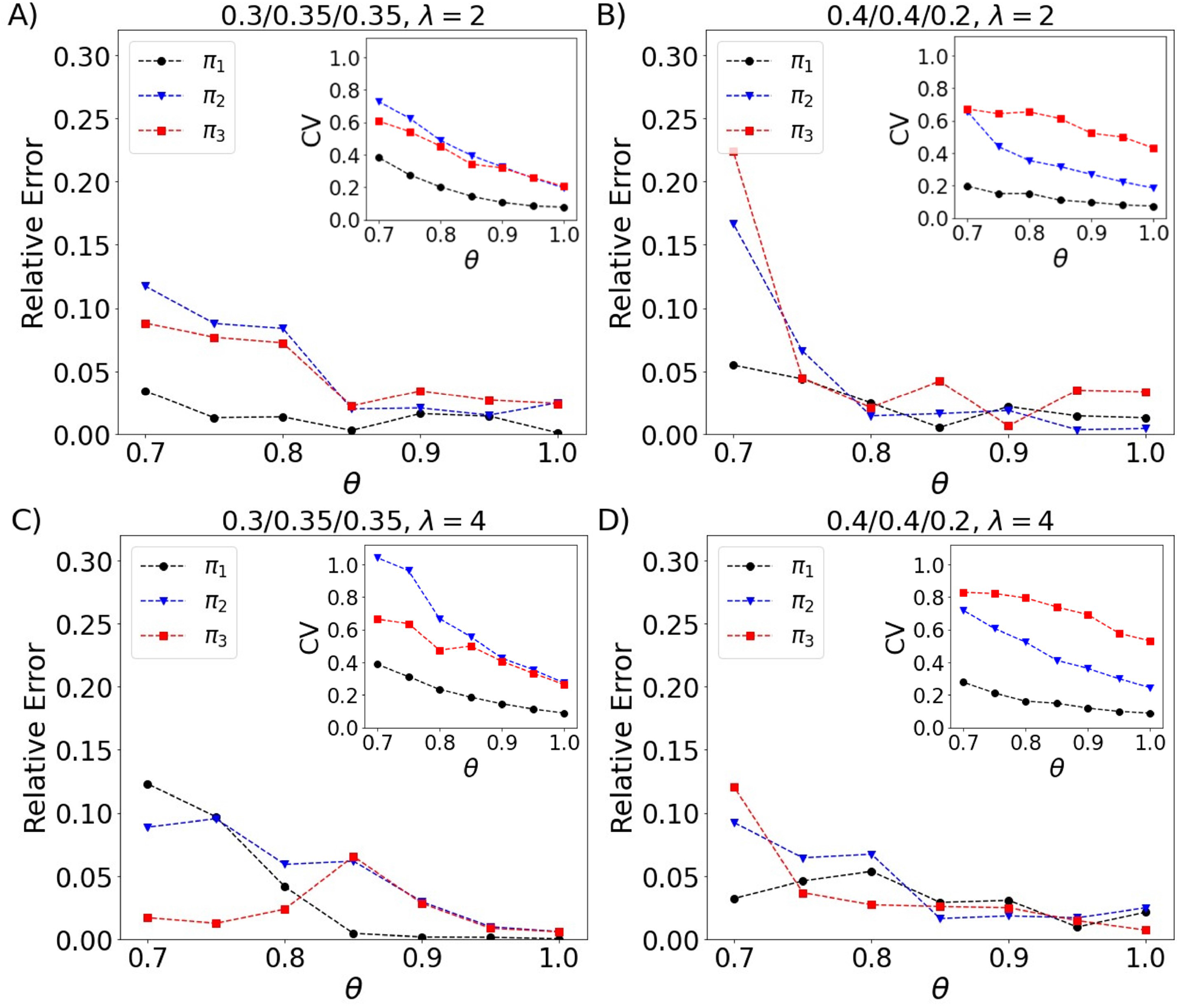
Effects of imperfect labeling for λ = 2 and 4 for M/D/T models of mixtures of 0.3/0.35/0.35 or 0.4/0.4/0.2 (simulations of 200 replicates, 1000 protomers each). The mean relative error as a function of *θ*. Inserts show the corresponding CV as a function of *θ*.

In conclusion, we have presented a machine learning approach to quantifying the monomer, dimer, and trimer fractions of an oligomeric protein from SMLM datasets. This is a significant extension of our previous method for SMLM based molecular counting that was built upon the exponentially distributed number of blinks emitted by many photoswitchable fluorophores. Both the novelty and power of this new method is that the algorithm is capable of learning the photophysical parameter λ needed to convert the observed number of blinks of a photoswitchable fluorophore to an estimate of molecule number. We found that the resulting statistical estimates of the protomer fractions are quite accurate when the molecules are well labeled, but the uncertainty in these estimates increase significantly with increasing labeling inefficiency. This would suggest that the method is more suited for SMLM measurements on proteins covalently labeled with organic dyes (e.g., dSTORM), where the labeling efficiency can be well over 90%, but is less relevant when working with fluorescent proteins (e.g., PALM) that often display much lower labeling efficiencies. While some of the uncertainty introduced by labeling inefficiency can be modulated by working with fluorophores that blink less, which is typically the case for fluorescent proteins, we found that reducing λ yielded only modest gains.

We end by stressing that, when applying this method, it is important to validate the underlying photophysical assumptions that the technique is built upon. Current organic dyes used for dSTORM are notoriously sensitive to their environment, and the intracellular milieu could alter the photophysical properties of the dyes in a heterogeneous manner so that the blinking statistics no longer follow a single-exponential distribution. Likewise, even fluorescent proteins, which are often thought to be shielded by the *β*-barrel, have shown environmental sensitivity [29]. This can only be checked by incorporating the proper controls, say by considering a purely monomeric or dimeric population, as should be done in any molecular counting experiment. Fortunately, new photoswitchable fluorophores are continually being developed for SMLM, both FPs that display more consistent and improved maturation and photoconversion efficiency and dyes whose blinking properties are better shielded from the environment [30, 31]. In conjunction with such technical advances, the advances presented here should make quantifying protein stoichiometry with SMLM more accessible and practical within a wider range of biological systems.

## Acknowledgments

We thank Alan Moses for fruitful discussions that stimulated this work. This research was funded by a Natural Sciences and Engineering Research Council (NSERC) Discovery Grant. A. Boonkird acknowledges support from a University of Toronto Excellence Award (UTEA).

