## Additional Supplemental Materials for "Quantitative Single-Molecule Imaging with Statistical Machine Learning"

#### Contents

**1 Simulation Details** **2**

**2 Experimental Details** **2**

**3 Convergence of EM Algorithm to $\lambda$** **3**

**4 Extracted Distribution of Blinks for Model M/D/T Mixtures** **4**

**5 Effects of Sample Size on the Performance of the AIC** **5**

**6 Effects of $\lambda$ on the Performance of the AIC** **6**

### 1 Simulation Details

The main assumption in our method is that the distribution of blinks per fluorophore follows a geometric distribution of the form

$$p(B|N = 1) = (1 - e^{-1/\lambda})e^{-(B-1)/\lambda}, \tag{S1}$$

where $\lambda$ is the characteristic number of blinks per fluorophore. As such, to simulate $N$ fluorophores, $N$ samples were drawn from this distribution to create a $N \times 1$ array, $\xi$, with the i-th entry denoting the number of blinks for the i-th fluorophore. Stoichiometric mixtures of monomers, dimers and trimers were created by randomly sampling with repetition from $\xi$. For dimers and trimers, the number of blinks for each fluorophore in the complex were summed. To model the detection efficiency of fluorophores, the number of flurophores for which we consider the blinks were determined by 'flipping a coin' with acceptance probability $\theta$ for each fluorophore chosen in a complex. Fluorophores that lost the coin flip did not have any of their blinks counted.

### 2 Experimental Details

Further details and all protocols concerning the single-molecule imaging of single dye DNA origami, used to experimentally validate our method, can be found in [1] . Briefly, cleaned glass coverslips were incubated in a 10% (v/v) poly-l-lysine (Sigma-Aldrich) solution overnight. The coverslips were then air dried, and covered for 25 minutes in a 5% v/v solution of gold nanoparticles (40nm diameter, Sigma-Aldrich) to be used as fiducial markers for post-processing drift correction. After washing with distilled water and air-drying, the coverslips were used to construct 20 $\mu$L flow chambers with double-sided tape on a microscope slides

The surface of the flow chamber was then functionalized with BSA-biotin and streptavidin to bind custom DNA origami rectangular grids with a streptavidin conjugated overhang at each corner of the grid and a single Alexa647-conjugated overhang at the centre of the grid. dSTORM imaging was carried out on a custom-built setup based on an Olympus IX-81 inverted microscope with a 60X, NA = 1.49 oil-immersion TIRF objective. The dSTORM imaging buffer consisted of a cocktail of an oxygen scavenging system (PCA/PCD at 13 mM and 50 nM respectively) in folding buffer ( 5mM Tris, 50mM NaCl, 1mM EDTA, and 15mM $MgCl_2$) and 10mM MEA to induce photoswtiching of Alexa647 fluorophores with 637nm irradiation (2.4 kW/cm$^2$.)

### 3 Convergence of EM Algorithm to $\lambda$

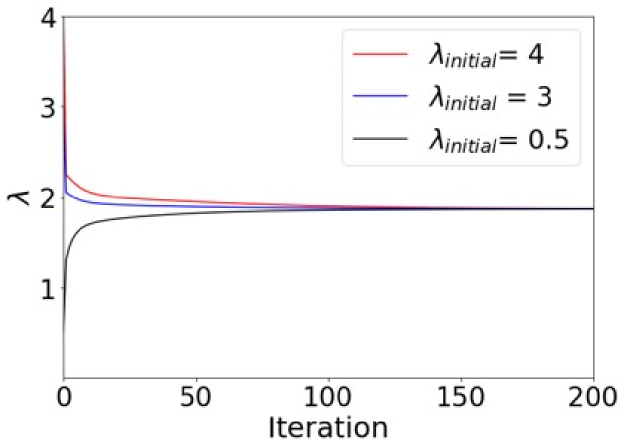


Figure S1: Convergence of the Expectation-Maximization (EM) algorithm to the expected value of the characteristic number of blinks, $\lambda = 2$ (simulated data). The initial guess $\lambda_{\text{init}}$ affects the number of iterations needed to achieve convergence to the expected value ($\lambda = 2$), but it does not affect the value to which the EM algorithm converges to.

### 4 Extracted Distribution of Blinks for Model M/D/T Mixtures

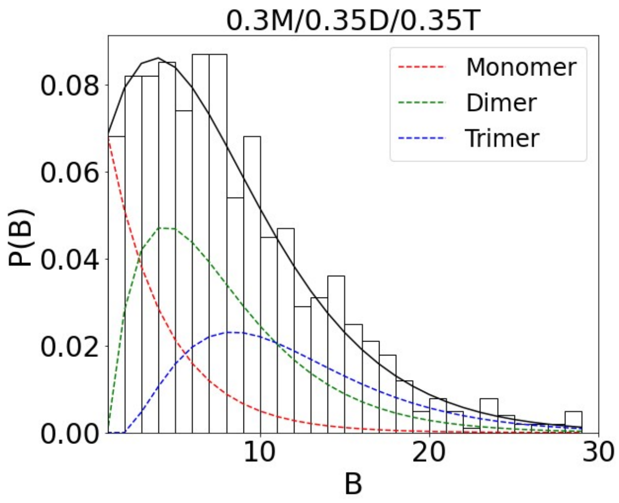


(a) 30/35/35

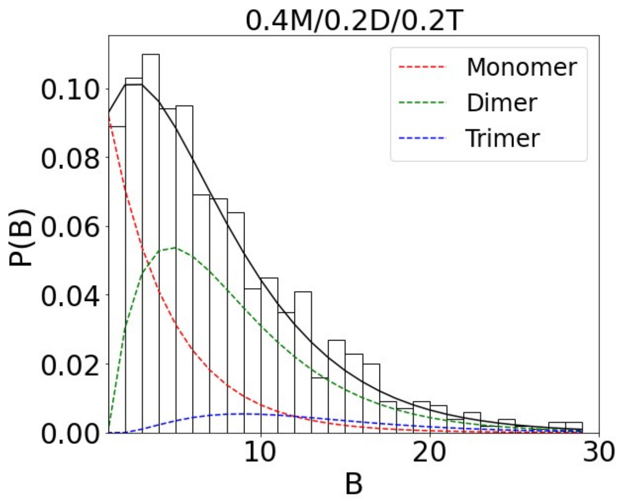


(b) 40/40/20

Figure S2: Explicit distributions for the two model stoichiometric mixtures decomposed from the experimental data. The contribution from each component found with the EM algorithm is shown with dashed lines (red: monomer, green: dimer, blue: trimer). The net distribution is shown with a solid black line.

### 5 Effects of Sample Size on the Performance of the AIC

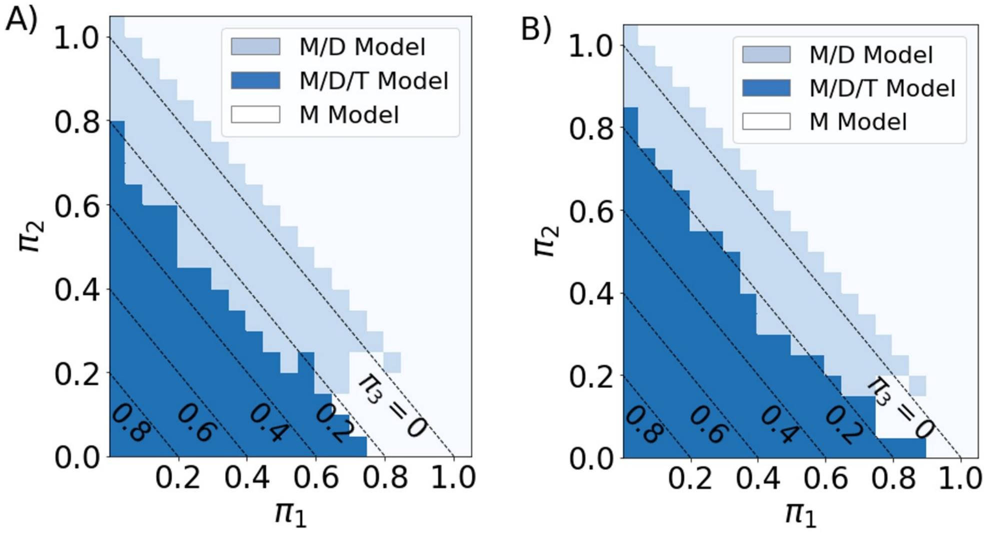


Figure S3: The accuracy of the AIC as a model selection criterion depends on the sample size. A) For a sample size of 1000 protein complexes (experimental data), which is a typical size of a data set in molecular counting experiments, the AIC is able to consistently select the correct mixture model (M/D/T) up to a trimer population of about $\pi_3 = 0.25$ and the corresponding range of values for $\pi_1$ and $\pi_2$. B) For a sample size of 3000 protein complexes (experimental data), the AIC is able to consistently select the correct mixture model (M/D/T) up to a relative trimer population of about $\pi_3 = 0.20$ and the corresponding range of values for $\pi_1$ and $\pi_2$. In the absence of a timer population, the AIC consistently selects the correct mixture model (M/D model). The AIC corrected for small sample sizes did not differ significantly from the non-corrected AIC for the sample sizes considered (data not shown).

### 6 Effects of $\lambda$ on the Performance of the AIC

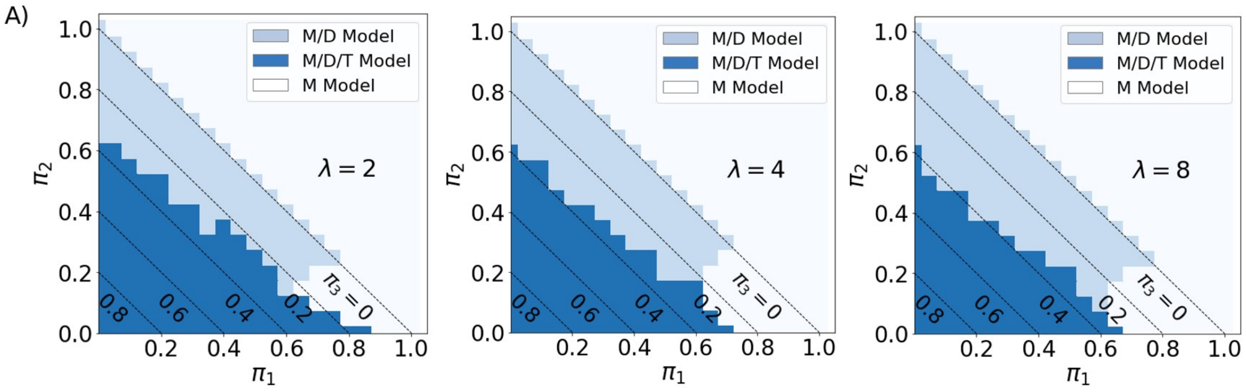


Figure S4: The accuracy of the AIC for model selection is sensitive to the magnitude of the characteristic number of blinks $\lambda$. For a sample size of 1000 simulated protein complexes, the range of accuracy of the AIC for model selection decreases for larger values of $\lambda$. A) For $\lambda = 2$, the AIC consistently selects the correct model (M/D/T) up to a trimer population of about $\pi_3 = 0.3$. B) For $\lambda = 4$, the AIC consistently selects the correct model (M/D/T) up to a trimer population of $\pi_3 = 0.35$. C) For $\lambda = 8$, the AIC consistently selects the correct model (M/D/T) up to a trimer population of about $\pi_3 = 0.4$. In the absence of a trimer population, the AIC consistently selects the correct mixture model (M/D model) for the range of $\lambda$ considered here.
